# Machine learning with biomedical ontologies

**DOI:** 10.1101/2020.05.07.082164

**Authors:** Maxat Kulmanov, Fatima Zohra Smaili, Xin Gao, Robert Hoehndorf

**Affiliations:** Computational Bioscience Research Center, Computer, Electrical and Mathematical Sciences & Engineering Division, King Abdullah University of Science and Technology, 4700 King Abdullah University of Science and Technology, Thuwal 23955-6900, Saudi Arabia

**Author notes:** **Author descriptions** **Maxat Kulmanov** is a postdoctoral researcher in computer science. His research interests include knowledge discovery and data integration using artificial intelligence and Semantic Web technologies in biology and biomedicine. **Fatima Zohra Smaili** is a doctoral student in computer science at King Abdullah University of Science and Technology. Her research focuses on ontology-based knowledge representation. **Xin Gao** is an Associate Professor in computer science, Acting Associate Director of the Computational Bioscience Research Center, and Lead of the Structural and Functional Bioinformatics Group at King Abdullah University of Science and Technology. His research focuses on bioinformatics and machine learning. **Robert Hoehndorf** is an Assistant Professor in computer science and principal investigator of the Bio-Ontology Research Group at King Abdullah University of Science and Technology. His research focuses on combining knowledge representation and machine learning in biology.

**Keywords:** machine learning, semantic similarity, ontology, knowledge representation, neuro-symbolic integration

## Abstract

Ontologies have long been employed in the life sciences to formally represent and reason over domain knowledge, and they are employed in almost every major biological database. Recently, ontologies are increasingly being used to provide background knowledge in similarity-based analysis and machine learning models. The methods employed to combine ontologies and machine learning are still novel and actively being developed. We provide an overview over the methods that use ontologies to compute similarity and incorporate them in machine learning methods; in particular, we outline how semantic similarity measures and ontology embeddings can exploit the background knowledge in biomedical ontologies, and how ontologies can provide constraints that improve machine learning models. The methods and experiments we describe are available as a set of executable notebooks, and we also provide a set of slides and additional resources at https://github.com/bio-ontology-research-group/machine-learning-with-ontologies.

**Key points:**

- Ontologies provide background knowledge that can be exploited in machine learning models.
- Ontology embeddings are structure-preserving maps from ontologies into vector spaces and provide an important method for utilizing ontologies in machine learning. Embeddings can preserve different structures in ontologies, including their graph structures, syntactic regularities, or their model-theoretic semantics.
- Axioms in ontologies, in particular those involving negation, can be used as constraints in optimization and machine learning to reduce the search space.

## 1 Introduction

Machine learning methods are now applied widely across life sciences to develop predictive models [1]. These models commonly solve an optimization problem, i.e., they perform search for an optimal solution to a function in a continuous or discrete space. Domain-specific knowledge can be used to constrain search and find optimal or near-optimal solutions faster, or to find better solutions; this observation has led Feigenbaum in 1977 to suggest that the power of Artificial Intelligence systems lies in the domain-specific knowledge they encode and are able to exploit, leading to the paradigm that “in the knowledge lies the power” [2].

In the life sciences, domain-specific knowledge is often encoded in ontologies and in the data- and knowledge-bases that use ontologies for annotation. Hundreds of ontologies have been developed, spanning almost all domains of biological and biomedical research. The main features biomedical ontologies provide are controlled vocabularies for characterizing biological phenomena, and as formalized knowledge bases that formally describe the phenomena within a domain and link them to other related domains. For example, phenotype ontologies are used for characterizing the phenotypes observed in a variety of model organism databases [3–6] as well as in human genetics [7, 8], and these ontologies provide a controlled set of classes, their labels, and definitions for the purpose of annotating the phenotypes observed in conditions recorded in databases. Moreover, phenotype ontologies are also interlinked with other ontologies through the use of formal axioms and can be used to relate the phenotype observations to biological functions, anatomical locations, developmental stages, or chemical substances [9, 10]. The majority of biomedical ontologies are formalized using the Web Ontology Language (OWL) [11], a language based on Description Logic (a decidable fragment of first order predicate logic). OWL comes with an explicit semantics that defines how statements made in OWL constrain the world in which these statements are interpreted – the “models” in which these statements are true.

The background knowledge contained in ontologies can be used in machine learning models for at least two different purposes: to expand or enrich features to be used, and to constraint the search for an optimal solution to a learning problem. Expanding or enriching features may make information available to a machine learning model that it would not be able to access without relying on ontologies. For example, linking phenotypes such as *cardiomyopathy* to the anatomical structures that are affected (i.e., the *heart*) can create novel and direct associations with other datasets that do not otherwise exist. In the example of *cardiomyopathy*, the link to *heart* as the anatomical structure can be used to relate the phenotype to gene expression in *heart* tissue or in cardiomyocytes; this constrain is given *a priori* through the axioms in phenotype, anatomy, and celltype ontology, and does not need to be discovered.

A second application of the knowledge in ontologies is to constrain the search for solutions to an optimization problem, and thereby finding a solution faster, finding a better solution, or finding a solution that is generalized better. One example of such a constraint is the true path rule that was originally proposed in the Gene Ontology [12], which states that if a gene product *G* has the potential to be involved in a process *P*_1_, and every process *P*_1_ is a part of another process *P*_2_, then *G* must also be involved in *P*_2_. This constraint is ‘hard’ in that it is not an empirical law or observation, but should hold in virtue of the definition of *P*_1_ and *P*_2_ – it is *impossible* for *G* to participate in *P*_1_ but not *P*_2_. For example, a gene product involved in *developmental cell growth* (`GO:0048588`) must be involved in *cell development* (`GO:0048468`) simply based on the definition of the two classes in the Gene Ontology.

It is now a challenge to identify general ways in which ontologies, and their underlying formalisms based on first order logic, can be combined with the modern machine learning models that are becoming so widespread. This challenge is not only one of research in Artificial Intelligence but exists throughout the life sciences due to the widespread use of ontologies and formalized knowledge bases in biology and biomedicine.

Here, we describe and review the state-of-the-art and recent advances in machine learning with biomedical ontologies. We use as a starting point in our review more traditional semanic similarity measures applied to ontologies; semantic similarity measures are a method from Artificial Intelligence that can determine the similarity between two or more entities using formalized background knowledge. We continue to introduce unsupervised, deep learning methods on ontologies that generate ‘embeddings’ for entities in ontologies, and we show that these embeddings can be used like semantic similarity measures while additionally allowing to overcome some of their limitations. Third, we highlight methods that use ontologies as constraints in optimization problems. We summarize the methods and tools we introduce in Table 1. We continue by introducing a novel benchmark dataset for machine learning with ontologies and demonstrating the methods we discuss on this dataset; we also make all experiments available as executable notebooks which can be adopted to other use cases. We finish by reviewing some of the main limitations and future research directions for the combination of ontologies and machine learning.

**Table 1:**
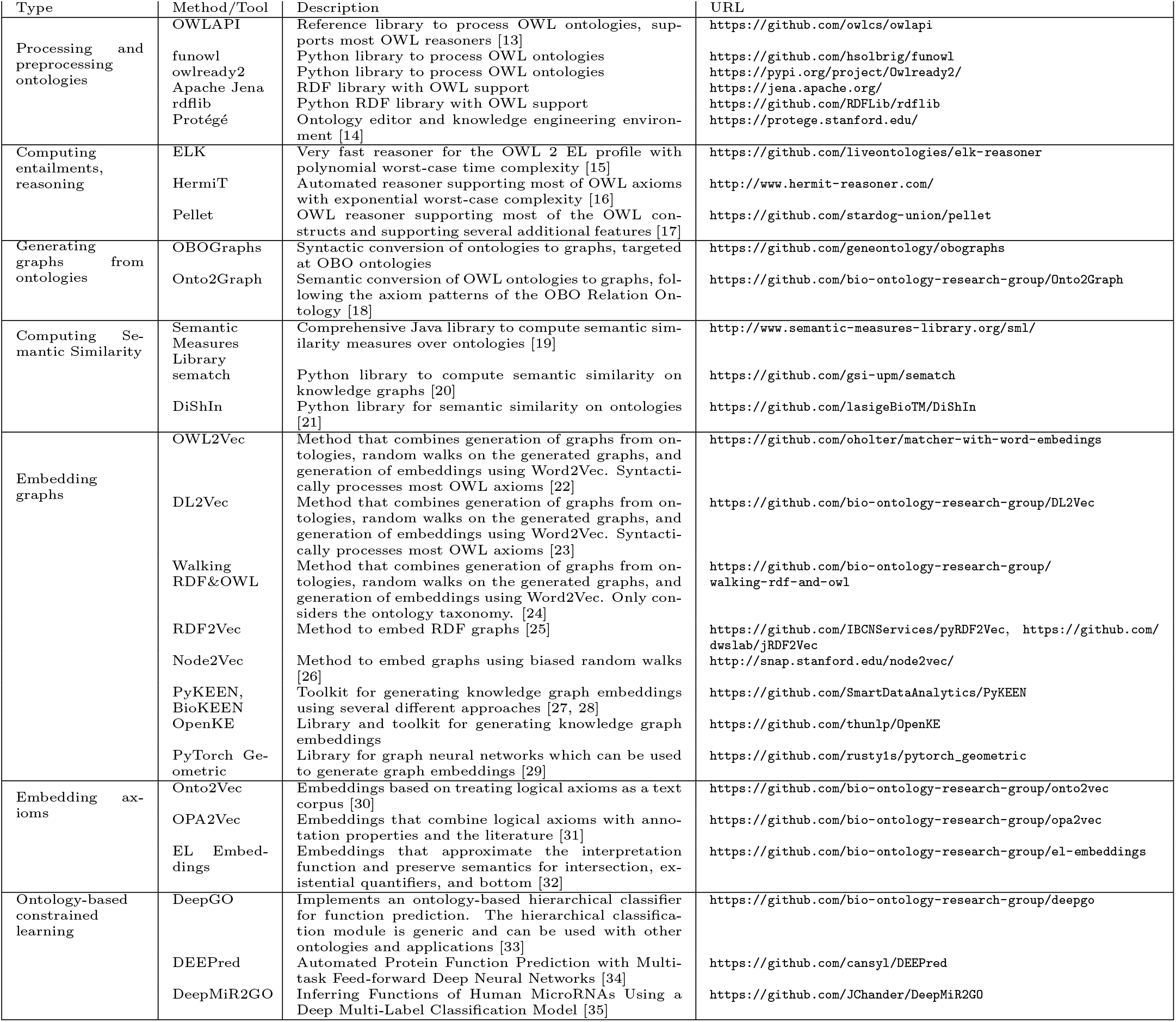
An overview of software tools and applications involved in machine learning with biomedical ontologies.

## 2 Axioms, graphs, and knowledge graphs

An ontology is an “explicit specification of a conceptualization of a domain” [36], i.e., an ontology is an artifact used to formally specify the intended meaning of a vocabulary within a domain. Ontologies contain domain knowledge, encoded in the form of axioms, natural language labels, synonyms, definitions, and other types of annotation properties. The majority of ontologies in the life sciences are encoded using the Web Ontology Language (OWL) [11], a language that is a part of the Semantic Web stack [37] and based on Description Logics [38]. Description Logics enable a formal, machine-readable description of the types of entities within a domain and the relations in which they stand [38]. Table 2 illustrates how the semantics of such a formal language (in this case, the Description Logic EL) is specified; syntactic constructs are assigned an interpretation in a mathematical structure that resembles a world in which these constructs are true. For example, *C* ⊑ *D* will be true in those structures in which all entities that are in the interpretation of *C* are also in the interpretation of *D*.

**Table 2:**
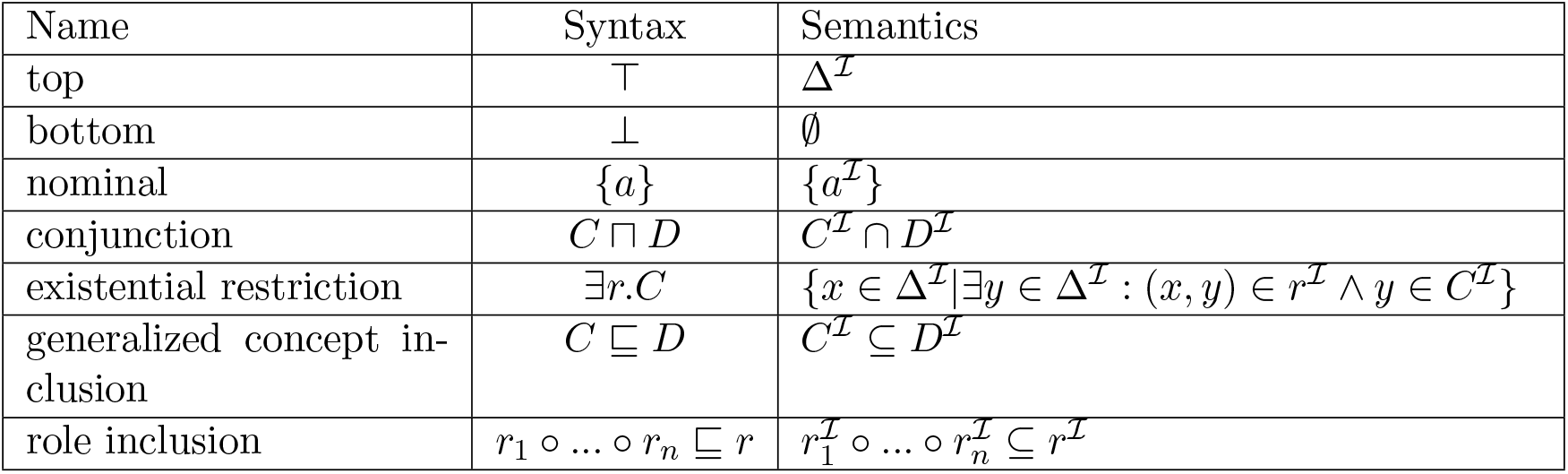
Description Logic EL

| Name | Syntax | Semantics |
| --- | --- | --- |
| top | $\top$ | $\Delta^{\mathcal{I}}$ |
| bottom | $\perp$ | ∅ |
| nominal | $\{a\}$ | $\{a^{\mathcal{I}}\}$ |
| conjunction | $C \sqcap D$ | $C^{\mathcal{I}} \cap D^{\mathcal{I}}$ |
| existential restriction | $\exists r.C$ | $\{x \in \Delta^{\mathcal{I}} \mid \exists y \in \Delta^{\mathcal{I}} : (x, y) \in r^{\mathcal{I}} \wedge y \in C^{\mathcal{I}}\}$ |
| generalized concept inclusion | $C \sqsubseteq D$ | $C^{\mathcal{I}} \subseteq D^{\mathcal{I}}$ |
| role inclusion | $r_1 \circ \dots \circ r_n \sqsubseteq r$ | $r_1^{\mathcal{I}} \circ \dots \circ r_n^{\mathcal{I}} \subseteq r^{\mathcal{I}}$ |

The semantics of logical languages gives rise to entailment; a statement *φ* is logically entailed by a set of statements *O* if all the structures in which all statements in *O* are true also make *φ* true. For example, the two statements {*C* ⊑ *D, D* ⊑ *E*} entail the statement *C* ⊑ *E*. The process of computing entailments – deduction or logical inference – plays a crucial role in using ontologies because it allows to automatically derive statements that are not explicitly asserted in a knowledge base, and can also be used to detect whether a set of statements is contradictory.

Many analysis methods that rely on ontologies, including machine learning methods and semantic similarity measures, rely on generating some form of graph structures from the axioms in an ontology. There are several ways in which axioms can be used to generate a graph structure, and many can be formulated as computing entailments. An important ontology for generating graphs from biomedical ontologies is the OBO Relation Ontology [39] which provides a set of axiom patterns that must hold true for two classes if an edge between them should be created. An axiom pattern is an axiom with variables for classes or individuals; *X* ⊑ *Y* is an axiom pattern in which *X* and *Y* are variables and if this statement is true for two classes *X* and *Y*, an edge labels is-a should be created between them: 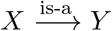. More complex axiom patterns involve quantifiers, such part-of as *X* ⊑ ∃part-of*.Y* which gives rise to the edge 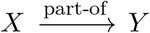. Axioms can also express disjointness between two classes such as *X* ⊓ *Y* ⊑ ⊥ based on which a disjoint edge can be created (*X* ↔ disjoint*Y*). To be generally applicable, these patterns must also be able to utilize entailments; for example, if *X* ⊑ ∃part-of*.Y* and *Y* ⊑ ∃part-of*.Z* are a part of an ontology, and the relation `part-of` is transitive (part-of ◦ part-of ⊆ part-of), then *X* part-of*.Z* would be entailed and consequently a `part-of` edge between *X* and *Z* part-of created (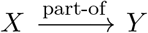). Depending on the algorithm that uses the graph, inferred edges can be added or not; for some relations, this amounts to adding their transitive closure, or, alternatively, transitive reduction [18].

The types and complexity of axiom patterns giving rise to edges is an active research area. For example, OWL2Vec [22] uses the set of transformation rules shown in Table 3 to transform syntactic axiom patterns into edges.

**Table 3:**
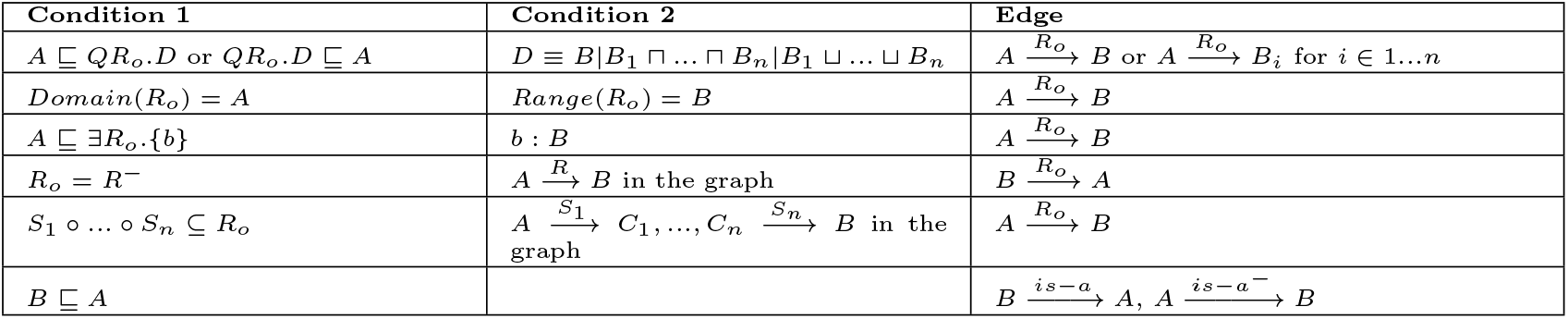
OWL2Vec rules for the projection of OWL axioms into an RDF graph. *Q* is any quantifier (∃, ∀, ≤ *n*, ≥ *n*, = *n*). *A*, *B*, *B*_*i*_ and *C*_*i*_ are named classes, *S*_*i*_, *R*_*o*_, and *R*^−^ are object properties, *b* an individual name.

The graphs generated from ontologies also interact with graph-based representations of data, in particular using the Resource Description Framework (RDF) [40]. Graphs in which nodes represent entities within a domain and edges represent the relations between the nodes are sometimes called *knowledge graphs*, and they correspond to a subset of the formalism underlying OWL in which only relations between individuals, and possibly certain axioms for relations, are considered. However, graph-based representations of the axioms in ontologies can also be considered knowledge graphs, in particular when both individuals and classes are included in the graph. For example, Figure 1 shows a graph in which interactions between proteins, the associations between proteins and their functions, and some axioms from the Gene Ontology are included. There are several ways in which such a graph could have been represented in OWL and then converted into such a graph representation using axiom patterns [41, 42]; for example, the edge between *MET* and *MAPK3* could arise from an axiom *MET* ⊑ ∃ activates *.MAPK*3 and the edge between *FOXP2* and *GO:0071625* from the axiom *FOXP*2 ⊑ ∃ hasFunction .GO:0071625. The dashed edge between *FOXP2* and *GO:0044708* is an edge that would be generated through entailment based on these axioms.

**Figure 1:**
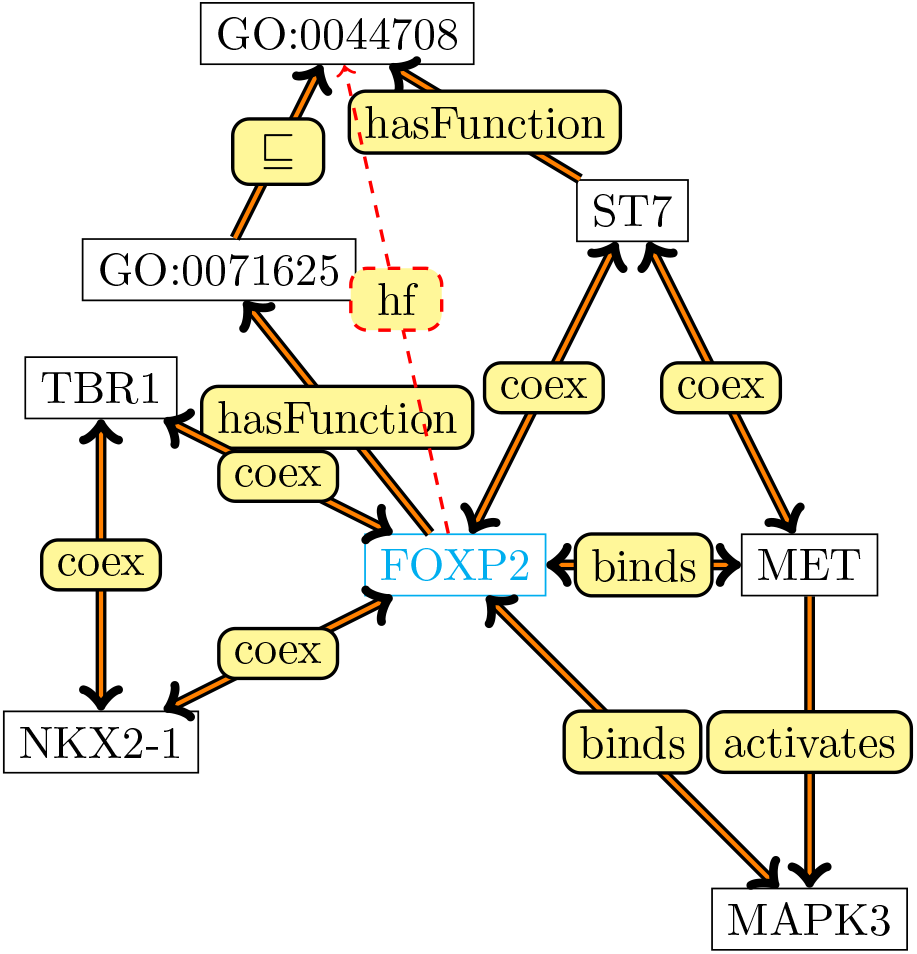
A knowledge graph centered around protein–protein interactions and functions of FOXP2.

## 3 Semantic similarity

Semantic similarity measures are widely used in biomedical domain. They are used to compare words, terms and ontology classes based on the background knowledge in the ontologies, annotations and large text corpora. Similarity measures are mostly hand-crafted and can be used as unsupervised classifiers for association prediction or as features in supervised learning models or a clustering algorithms. They can be applied to a variety of tasks such as predicting protein–protein interactions [43], gene–disease associations [44, 45], diagnosing patients [46, 47], determining sequence similarity [48], or evaluating computational methods which predict ontology class annotations [49].

We can compute semantic similarity measures between classes, class instances and annotated entities. A function *sim* : *D* × *D* is a similarity on a domain *D* if it is non-negative (*sim*(*x, y*) ≤ 0), symmetric (*sim*(*x, y*) = *sim*(*y, x*)), and if self-similarity yields the highest similarity values within the domain (*sim*(*x, x*) = max_*D*_), or – as a weaker version – if self-similarity is higher than similarity to any other domain entity (*sim*(*x, x*) *> sim*(*x, y*)).

A simple similarity measure, *sim_Rada_*, can be based on the shortest path between two nodes in the graph [50]. It can be defined as:

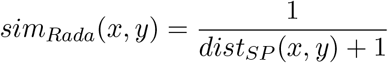

This similarity measure is useful when edges in a graph correspond mostly uniformly to some kind of semantic distance. However, when comparing ontology classes, edges represent axioms involving two classes which may not correspond to this assumption. For example, `is-a` edges order classes from general to more specific, such as in the ontology in Figure 2a. In this figure, *sim_Rada_*(*Color, Shape*) will have the same value as *sim_Rada_*(*Red, Green*) since these two classes have the same distance in the graph. However, in many applications *Red* and *Green* should be more similar than *Color* and *Shape* because they are both colors. In this case, distance based similarities might not be very intuitive and a measure of node specificity needs to be considered.

**Figure 2:**
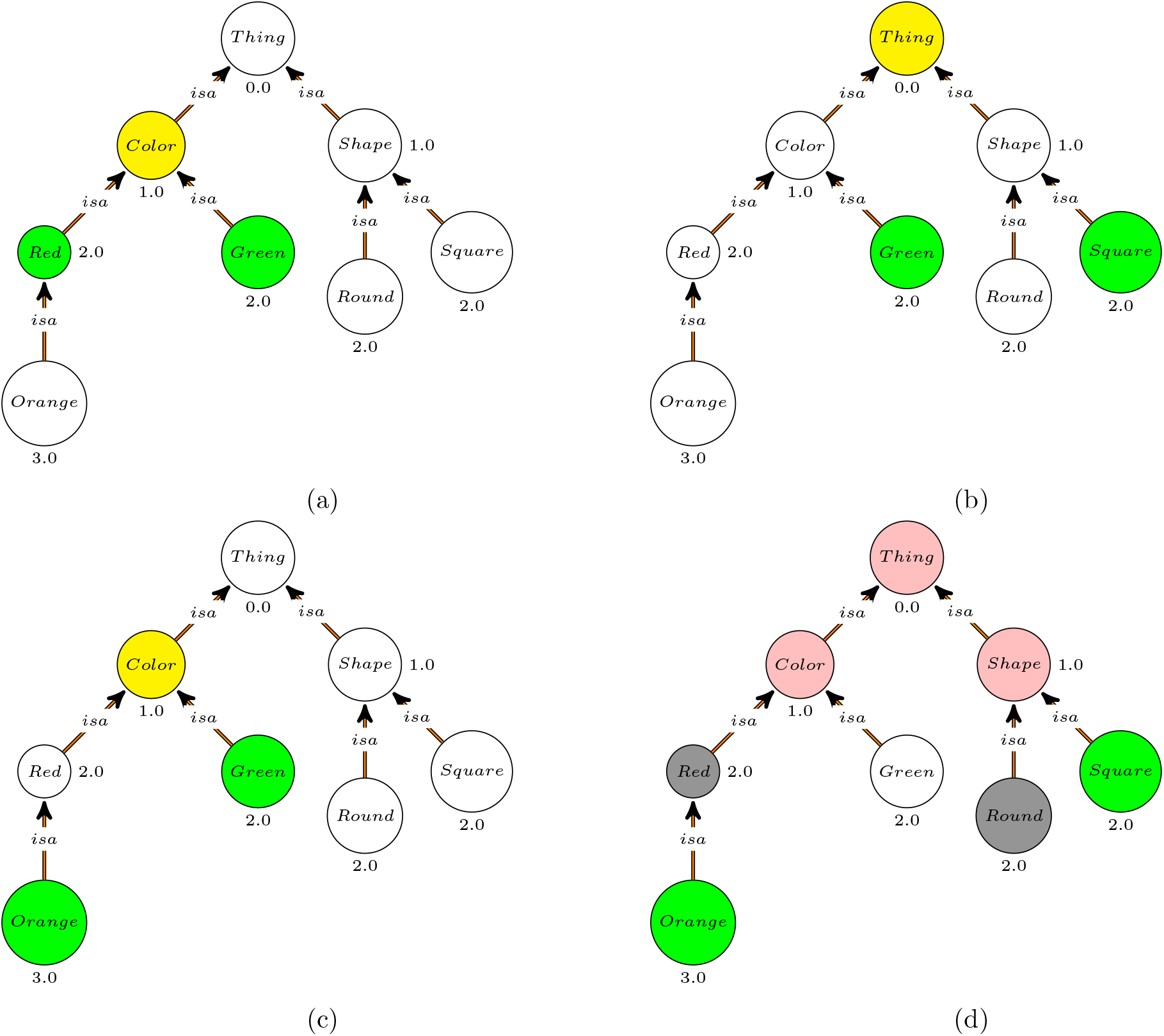
A fragment of the PATO ontology focusing on colors and shapes. Numbers near classes indicate specificity and information content of the classes.

There are many ways to compute class specificity. For instance, we can consider specificity as a function of the depth, number of children, or the information content of a class. Formally, class specificity is a function *σ* : *C* ↦ ℝ which meets the condition that for all *x, y* ∈ *C*, if *x* ⊑ *y* then *σ*(*x*) ≥ *σ*(*y*) [51]. The specificity measure can be defined using only the classes within an ontology (such as measures that consider the number of super-classes a class has, or the distance of a class to the root), or using information such as the number of instances of a class, or the number of annotations of a class within a database.

One of the most widely used methods to determine class specificity is the Resnik measure [52], which defines the specificity of a class as its information content:

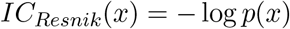

Where

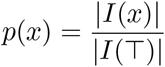

and *I*(*x*) is the set of instances of *x* (or the set of annotations of a class within a database).

A large number of semantic similarity measures have been developed [51]. Pairwise similarity measures compute the similarity value between two classes. Examples of pairwise similarities used in the biomedical field include Resnik’s [52], Lin’s [53], Jiang & Conrath’s [54] and Schlicker’s [44] similarity measures. Many of these measures are variations of the Resnik measure which defines the similarity between classes *x* and *y* as the information content of their *most informative common ancestor (MICA)*:

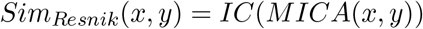

In the example in Figure 2a, *Sim*_*Resnik*_(*Red, Green*) is equal to 1.0 and *Sim*_*Resnik*_(*Color, Shape*) is equal to 0.0 although they have the same distance. The downside of this similarity measure is that it does not take into account the specificity of the compared classes and all classes under the same MICA will have the same similarity value. For instance, in Figure 2b *Sim*_*Resnik*_(*Green, Square*) is equal to 0.0 which is the same as *Sim*_*Resnik*_(*Color, Shape*) and in Figure 2c *Sim*_*Resnik*_(*Red, Green*) and *Sim*_*Resnik*_(*Orange, Green*) are both equal to 1.0. To solve this issue, Lin’s measure [53] also considers information content of the compared classes:

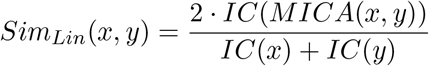

With this measure, *Sim*_*Lin*_(*Red, Green*) is equal to 0.5 whereas *Sim*_*Lin*_(*Orange, Green*) is equal to 0.4 which is more intuitive.

When comparing two instances of ontology classes, or two entities annotated with classes in an ontology, we usually need to compare sets of classes. For example, we would have to compute the similarity of the set of all Gene Ontology annotations of one protein with the set of all Gene Ontology annotations of a second protein. There are two ways of determining the similarity between two sets of classes *A* and *B*. First, we can compute the pairwise similarities between all pairs of classes (*a, b*) such that *a* ∈ *A* and *b* ∈ *B*, and then combine similarity values according to some combination strategy (such as computing the average). Second, we can directly define a similarity measure between the two sets *A* and *B* using a set similarity measure. For instance, we can use the Jaccard index between the two sets:

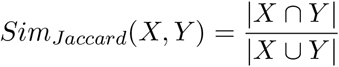

To make this a semantic similarity, we would at least close each of the sets *X* and *Y* with respect to superclass axioms, i.e., if *C* ⊑ *D* and *C* ∈ *X* then *D* ∈ *X*. Figure 2d depicts the propagation of ontology classes for computing the similarity between a square-and-orange thing and a round-and-red thing. Set similarity can also incorporate class specificity, such as the weighted Jaccard index in the SimGIC [55] measure.

Semantic similarity measures have a variety of applications and a large number of software packages have been developed to ease their use. One prominent example is the Semantic Measures Library [19] which is a comprehensive Java library that allows to compute hundreds of different semantic similarity measures.

A common problem of semantic similarity measures is that it is difficult to choose one for a particular application. Similarity measures behave differently depending on their applications. For example, predicting protein–protein interactions may result in different performance [55, 56] with similarity measures depending on the organisms. They are not immune to biases in data and different similarities may react to the biases differently [57]. Furthermore, they are hand-crafted measures that are not able to adapt to the underlying data or application.

## 4 Embedding ontologies

Another option to define similarity measures on ontologies is through the use of embeddings. An embedding is a structure-preserving map from one mathematical structure to another. We can use embeddings to project the elements of one structure into a second one. The idea behind using embeddings is that the second structure may enable different or additional operations which are not possible in the first structure. For example, if we take ontologies or graphs that are discrete entities and map them into a continuous space (or real-valued vector space), we can apply machine learning or continuous optimization algorithms which operate on continuous data; there are also ‘natural’ similarity measures between real-valued vectors such as the cosine similarity or other distance measures and metrics.

In the context of machine learning in most cases, we aim to embed ontologies within real-valued vector spaces. The key question when embedding ontologies is which structure (of the ontology) to preserve within ℝ^*n*^ and under which operations in ℝ^*n*^ this structure is preserved. We classify approaches of embedding ontologies in three main types based on what aspect of the ontologies is preserved in ℝ^*n*^. First, there are *graph-based* approaches which treat ontologies as graphs similar to how ontologies are treated by many semantic similarity measures, and the embeddings preserve this graph structure within ℝ^*n*^. Second, *syntactic* approaches treat axioms similar to “sentences” and preserve syntactic regularities (such as frequencies of co-occurrences) in ℝ^*n*^. Third, we consider *model-theoretic* approaches which preserve model-theoretic properties within ℝ^*n*^ as a part of the embedding.

### 4.1 Graph-based ontology embeddings

Graph-based embedding methods preserve a graph structure within ℝ^*n*^. One form of graph embeddings is based on random walks. In these methods, graphs are generated from ontologies using the methods we described, then random walks are used to explore the neighborhood of each node in the graph, and finally the set of walks is used as the basis of the embeddings.

One of the first methods for learning graph embeddings through random walks was DeepWalk [58] which generates a ‘corpus’ of sentences (i.e., sequences of nodes in the graph) through random walks starting from each node in the graph, and then applies Word2Vec on the resulting corpus to obtain embedding vectors; the embeddings generated by Word2Vec preserve co-occurrence relations within a context window. DeepWalk can also be extended to include labeled edges [22, 24, 25].

For example, for the graph in Figure 1, the random walks can generate sentences such as

- FOXP2 cooex ST7 hasFunction GO:0044708 …
- FOXP2 hasFunction GO:0071625 is-a GO:0044708 …

and Word2Vec will then embed each node and edge label while preserving co-occurrence relations within this corpus [60]. Node2Vec [26] is a modified model that does explore the original graph through ‘biased’ random walks and therefore can force walks to remain within a certain distance of the origin node or explore further away.

Random walks have long been used as a model that simulates diffusion of information within a network [61–63] and can be used to identify and score node importance. In graph embeddings, these walks explore node neighborhood and generate a ‘linear’ representation (i.e., sequences of symbols) in which nodes that are reached more often also occur more often (and co-occur more often with the original node). Word2Vec, as a model that embeds sequences of symbols while maintaining this co-occurrence, generates embeddings that maintain this syntactic structure within the walks, and therefore aspects of the graph structure as well. Furthermore, some of the semantics of the axioms in the ontology can be encoded as constraints on the random walks or encoded in the graph; for example, symmetry can be modeled as a bi-directional edge, disjointness as a ‘barrier’ preventing a walk’s transition, etc. It is obvious that the graph that is generated from the ontology axioms, and the information it captures, is crucial for generating useful embeddings; this is also an active research area [22].

Translational embeddings methods are a family of representation learning methods on knowledge graphs which model relations in the knowledge graph as translation operations between graph node embeddings. The methods have been successfully applied for several tasks such as link prediction, knowledge-graph completion and others. The methods represent knowledge graphs as a set of edges (*s, p, o*) (triples) and define a translation operation which translates *f*_*η*_(*s*) to *f*_*η*_(*o*) depending on the relation *p*. Here, *f*_*η*_ is a graph embedding.

TransE [64] was the first translational embedding method. It uses a vector representation for relations that have the same dimensions as vectors representing nodes, and defines the translation operation as the addition of the relation vector to the node vector:

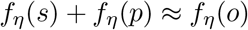

and further defines a scoring function for an edge based on the translation operation:

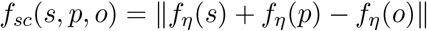

Then, it minimizes the following loss function to learn *f*_*η*_:

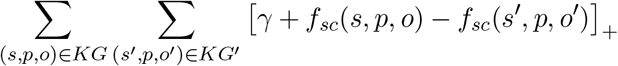

where *KG*′ is a set of negative or corrupted triples that are not in the graph, [*x*] _+_ indicates the positive part of *x*, and *γ* is a hyperparameter. This model can only accurately represent one-to-one relations and it is not suitable for one-to-many and many-to-many relations; in graphs generated from ontologies, even when focusing only on the subclass hierarchy, there are many such relations. Furthermore, TransE does not support transitive, symmetric or reflexive relations which are all important for faithfully embedding ontologies.

Many TransE successors have been developed to overcome the original model’s limitations. For example, TransH [65] extended TransE by moving the translation operation to a relation-specific hyperplane. TransH represents each relation by two embedding vectors, the norm vector of the hyperplane (denoted as a function *w*_*η*_) and a translation vector (denoted as a function *d*_*η*_). The scoring function is then defined as:

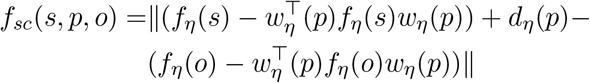

With an additional vector, TransH performs translation operation on an augmented hyperplane and can therefore model one-to-many and many-to-many relations better than TransE. There are also many other models with various advantages and disadvantages [66, 67].

Translational embeddings are able to explicitly capture the graph structure and preserve some interpretability through the use of vector operations; however, they cannot easily capture even simple axioms such as transitivity, symmetry, or reflexivity of relations. Furthermore, any graph-based method will focus on a small set of graph patterns and lose some information about the ontology axioms, while many ontological axioms such as disjointness and axioms involving combinations of different logical operators cannot be fully converted to a graph.

### 4.2 Syntactic approaches

Ontologies provide a structured representation of biological knowledge in the form of logical axioms, and not all the axioms in an ontology can be represented naturally in a graph; this limits the ability of these methods to utilize all information encoded in the ontology. Syntactic embeddings embed ontologies in ℝ^*n*^ considering only the set of axioms without creating an intermediate graph-based representation.

Onto2Vec [30] is a method that generates embeddings for ontology classes and instances taking into account the logical axioms that define the semantics of ontology classes. Onto2Vec takes an ontology *O* as input, uses a reasoner to infer additional logical axioms, mainly subclass axioms between named classes; it then treats each asserted or inferred axiom as a sentence and embeds the set of axioms using the Word2Vec language model. This allows Onto2Vec to embed ontologies directly based on their axioms while considering all axiom types, no matter how complex they are.

OPA2Vec [31] extends Onto2Vec to not only include logical axioms but annotation properties as well. Annotation properties in biomedical ontologies provide labels, synonyms, definitions, and other types of information about classes and instances in ontologies. OPA2Vec combines the corpus generated from the asserted and inferred logical axioms in Onto2Vec with a corpus generated from all or selected annotation properties. For example, from the annotation assertion that an OWL class *C* has a label *L* (using the `rdfs:label` annotation property in the OWL annotation axiom), OPA2Vec generates the statement `C rdfs:label L`, using the complete identifier for *C* and `rdfs:label`, and expressing *L* as a string literal; for instance, the annotation assertion of the class *Nuclear periphery* (GO:0034399) and its label is expressed as the sentence <http://purl.obolibrary.org/obo/GO0034399> <http://www.w3.org/2000/01/rdf-schema#label> `nuclear periphery`. The identifier for the class *C* occurs within the ontology axioms and obtains parts of its meaning through the axioms; to ensure that the natural language terms used in the annotation properties have their ‘natural’ meaning as used in biomedical texts, OPA2Vec uses transfer learning and pre-trains a Word2Vec language model on biomedical literature texts, and then tunes it to generate the embeddings from the axioms plus annotation properties.

### 4.3 Model-theoretic or semantic

None of the embedding methods discussed so far are ‘semantic’ in the sense that they use the semantics of the underlying logic (as, for example, shown in Table 2). Instead, the models are based on syntactic co-occurences or preserving certain graph properties. However, the main advantage of using languages with an explicit semantics is that it provides constrains on how symbols should be interpreted.

EL Embeddings [32] aim to embed ontologies by mapping the symbols in the ontology into one specific interpretation, i.e., the embedding is identical to, or approximates, the interpretation function 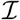 in Table 2. EL Embeedings find an embedding that maps the class, relation, and instance symbols Σ(*O*) into ℝ^*n*^, *f*_*e*_ : Σ(*O*) ↦ ℝ^*n*^ such that *f*_*e*_(Σ(*O*)) is a model of *O* (*f*_*e*_(Σ(*O*)) ╞ *O*). Such an embedding yields a faithful representation of logical operators and quantifiers.

Formally, EL Embeddings embed classes as *n*-balls in *n*-dimensional space and relations as *n*-dimensional vectors. The correspondence with the semantics of the axioms in the ontology is established by setting the domain of discourse to ℝ^*n*^ and the following condition: for all classes *C* ∈ Σ(*T*) and relations *r* ∈ Σ(*T*) it defines 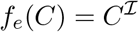 :

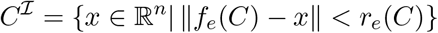

where *r*_*e*_(*C*) is the radius of the *n*-ball that corresponds to *C*, and 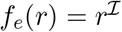 :

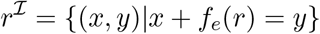

The latter condition is similar to the TransE translation operation only applied to instances.

The embeddings are generated through optimization using a set of loss functions that correspond to different normal forms of the axioms in ontologies. Using these embeddings it is possible to approximate the intended semantics of the language within the embedding space. In particular, it can be shown that if the loss can be reduced to zero, the resulting embedding corresponds to a model of the ontology [32]. Similar approaches to EL Embeddings are also investigated for querying knowledge graphs using logic formulas[68].

### 4.4 Using embeddings as semantic similarity measures and for machine learning

Embeddings can generate distributed representations of the symbols in ontologies while preserving syntactic or semantic properties. These representations – vectors in ℝ^*n*^ can be visualized using dimensionality reduction techniques such as principal component analysis or tSNE [69]. They can also be used to compute similarity using any kind of similarity or distance measure applicable to real-valued vectors, in particular the cosine similarity or the Euclidean distance. However, these similarity measures are still generic and do not adapt to particular applications.

One of the most useful applications of ontology embeddings is as a part of machine learning models in which either a single embedding is used as input or multiple embeddings are used as input. Single embeddings can be used in classification and related tasks while multiple embeddings are often used to predict relations between the entities which were embedded [67]. Such an application is similar to using a semantic similarity measure with the key difference that the actual function that computes the similarity can be ‘trained’ in a task-specific way using supervised learning. For example, if the similarity between two protein embeddings is supposed to be a measure of whether or not they interact, using a set of proteins that interact can be used to design a customized, task-specific function that predicts, given two embeddings as an input, whether the proteins they represent should interact. Many neural network architectures and other machine learning models can be used for this task, but architectures that are used for similarity learning, such as Siamese neural networks, seem to perform well in practice [23].

## 5 Ontologies as constraints

Ontologies embeddings are a useful technique to make information in ontologies available as background knowledge to define similarity measures or use as features in machine learning models. In these cases, ontologies are used as the *input* of a similarity function or a model. However, ontologies can also be used as an *output* of a machine learning model and the axioms in the ontology used to constrain the output of a function, such as in the case when determining if the predictions of a machine learning model are consistent with the axioms in the ontology.

Ontologies are used as structured output in many domains in which the primary task is to predict whether some entity has a relation with one or more ontology classes, such as predicting genotype–phenotype relations (using phenotype ontologies as output), predicting gene–disease or drug–disease associations (using disease ontologies as output), or predicting protein functions (using the Gene Ontology as output). At the very least, these tasks need to satisfy the hierarchical constraints imposed by the ontologies in the output space: if an entity *e* is predicted to be associated with a class *C*, and that class *C* is a subclass of *D*, then *e* must also be associated with *D*. Similar constraints arise from other axioms in the ontology.

In general, there are at least five different approaches to using hierarchical relationships as constraints in classification models: flat, local per node, local per parent, local per level, and global hierarchical classification [70]. Flat classification is when the hierarchical constraints are not used during the prediction or training and the classification is done only using individual classes, and the consistency with the hierarchical constraints is enforced by propagating scores along the hierarchy only after predictions are made. This approach employs the constraints imposed by the ontology independently from the training or prediction process. In a local per node setting, a binary classifier is built for every class and predictions are made starting from the most general classes first and then moving to more specific ones, and stopping the prediction process once classes are predicted as negative. In a local per parent and local per level setting, multi-class classifiers are used for children classes of a parent or classes at the same level, respectively. Similarly to local per node classifiers, the prediction is performed in a step-wise manner from the most general class to more specific ones, and terminated once predictions are negative. The main drawback of local classifiers is that all classification models are trained independently from each other, and during the prediction process errors will propagate from general classes to more specific ones. Global hierarchical classifiers include the hierarchical constraints during training of a machine learning model, either as soft constraints or hard constraints, and also during prediction so that the output labels are forced to be consistent with the ontology axioms. The advantages of these models are that they take the semantics into account during training and therefore potentially reduce the search space; and that they can exploit dependencies between classes during training and prediction; the disadvantage is often the increased complexity of these classifiers [70].

While hierarchical machine learning models are used across many different application domains, life science ontologies are standing out with their large size and complex set of axioms; it is no surprise that constrained optimization methods applicable to large ontologies have emerged from research in bioinformatics. In particular prediction of functions and phenotypes benefits from machine learning models that are constrained by ontologies, and the established evaluation measures for predictive performance of models in these domains are also ontology-based [71]. A large number of ontology-based prediction models have been developed in these domains, using hierarchical top-down phenotype prediction [72], using structured support vector machines for predicting functions [73] and phenotypes [74], and several methods that incorporate ontology-based constraints in artificial neural networks [35, 75, 76].

## 6 Use case and application

Ontologies are used in almost every major biological database. There are more than 800 ontologies in ontology repositories such as BioPortal [77] which are used to describe different biological and biomedical entities. Consequently, ontologies play a role in many different biomedical machine learning tasks such as genotype–phenotype association prediction [46, 47], protein function prediction [49], drug–target prediction [78, 79], protein–protein interaction prediction [21, 48, 56], gene–disease association prediction [80], and many others.

Here, we evaluate ontology embedding methods on the task of predicting interactions between proteins based on the hypothesis that functionally related proteins are more likely to interact. We demonstrate how different ontology embedding methods can be applied, and we provide Jupyter Notebooks for all our experiments at https://github.com/bio-ontology-research-group/machine-learning-with-ontologies.

Proteins do not function in isolation, and many biological processes and functions are regulated by multiple proteins and their interactions. Databases such as String [81] collect information about protein–protein interactions (PPIs) from different sources with experimental evidence as well as PPIs that are computationally inferred and automatically assigned, and the functions of proteins are described using the Gene Ontology [12].

We created two PPI datasets, one for interactions occurring in humans and one for yeast, based on data from the String database [81]. We filtered out interactions with a confidence score less than 700 to retain only high confidence interactions. Table 4 provides the total number of proteins and interactions in each dataset. We split the two datasets consisting of interaction pairs into train and test sets, with a ratio of 80% and 20%, respectively, and we used 20% of the training set as a validation set. We used these two datasets as benchmark sets for evaluating ontology embedding and semantic similarity methods, and we made the datasets with documentation publicly available for download and provided the links in our public repository so that anybody can use the same data to benchmark and compare ontology-based prediction methods. The training and validation sets should be used to train and tune model parameters and select the best models, while the evaluation results and comparisons should be reported using the test set.

**Table 4:** The total number of proteins and number of unique interaction pairs in training, testing, and validation datasets

| Organism | Proteins | Total | Training | Validation | Testing |
| --- | --- | --- | --- | --- | --- |
| Yeast | 6,157 | 119,051 | 76,193 | 19,048 | 23,810 |
| Human | 17,185 | 420,534 | 269,143 | 67,285 | 84,106 |

We predicted PPIs based on the associations of proteins with their functions and cellular locations represented in the GO, known interactions between proteins, and the information contained in the GO. One key question is how to represent these three types of knowledge as axioms in an ontology or knowledge base. We adopted a representation scheme in which all entities (proteins, functions, cellular locations) are classes and the relations between the entities are expressed as axioms [41, 42]. Specifically, if there is an interaction between proteins *P*_1_ and *P*_2_, we asserted the axioms *P*_1_ ⊑ ∃interacts-with*.P*_2_ and *P*_2_ ⊑ ∃interacts-with*.P*_1_; if protein *P* is associated with a GO class *C*, we asserted the axiom *P* ⊑ ∃has-function*.C*. We combined this set of axioms with the GO (released on 22 February 2020) to form our knowledge base.

For graph-based embedding methods, we generated a graph by creating an edge for existential restrictions in subclass axioms: if *X* ⊑ ∃*R.Y* is an (asserted) axiom in the knowledge base (consisting of GO together with the axioms we added), we created nodes *X* and *Y*, and an edge from *X* to *Y* labeled *R*. For the Onto2Vec and OPA2Vec embedding methods, we only inputted GO together with the set of protein-to-GO associations. The Jupyter Notebook *data.ipynb* in our repository provides source code to generate the datasets, the splits, and the input files for the different embedding methods.

We then generated ontology embeddings using EL Embeddings, Onto2Vec, OPA2Vec, and used the generated graph to produce embeddings through random walks, biased random walks (Node2Vec), and TransE. We then use these embeddings, as well as two semantic similarity measures (Resnik’s and Lin’s), to predict protein–protein interactions. For embeddings based on random walks, Onto2Vec, and OPA2Vec, we use cosine similarity to compute the pairwise similarity of all pairs of proteins in our dataset, including the proteins in the training, test, and validation sets; for TransE embeddings and EL Embeddings we use the prediction function for `interacts-with` edges and compute a prediction score for all pairs of proteins in our dataset; and we use the semantic similarity measures to compute the similarity between all pairs of proteins.

In the evaluation, for each protein *p* we rank all other proteins *p*_*i*_ based on their similarity to *p*. We then consider positives as pairs (*p, p*_*k*_) which are PPIs included in our test set, and we report hits (recall) at ranks 10 and 100, mean rank at which the PPIs are found, and the ROCAUC (using micro-averages per protein). Results are separated in *Raw* and *Filtered*; *Raw* results evaluate all pairs of proteins while *Filtered* results evaluate all pairs of proteins except the pairs that are included in the training or validation sets. *Filtered* results are usually better since training pairs are not considered in the evaluation. We made Jupyter Notebooks available for all our experiments, and Table 5 summarizes the results for yeast and Table 6 for human; all results in these tables can be reproduced using the Jupyter Notebooks.

**Table 5:** Prediction performance for yeast protein–protein interactions.

| Method | Raw Hits@10 | Filtered Hits@10 | Raw Hits@10 | Filtered Hits@100 | Raw Mean Rank | Filtered Mean Rank | Raw AUC | Filtered AUC |
| --- | --- | --- | --- | --- | --- | --- | --- | --- |
| TransE | 0.06 | 0.13 | 0.32 | 0.40 | 1125.4 | 1074.8 | 0.82 | 0.83 |
| SimResnik | **0.09** | **0.17** | 0.38 | 0.48 | 757.8 | 706.9 | 0.86 | 0.87 |
| SimLin | 0.08 | 0.15 | 0.33 | 0.41 | 875.4 | 824.5 | 0.84 | 0.85 |
| EL Em-beddings | 0.08 | **0.17** | **0.44** | **0.62** | **451.29** | **394.04** | **0.92** | **0.93** |
| Onto2Vec | 0.08 | 0.15 | 0.35 | 0.48 | 641.1 | 587.9 | 0.79 | 0.80 |
| OPA2Vec | 0.06 | 0.13 | 0.39 | 0.58 | 523.3 | 466.6 | 0.87 | 0.88 |
| Random walk | 0.06 | 0.13 | 0.31 | 0.40 | 612.6 | 587.4 | 0.87 | 0.88 |
| Node2Vec | 0.07 | 0.15 | 0.36 | 0.46 | 589.1 | 522.4 | 0.87 | 0.88 |

**Table 6:**
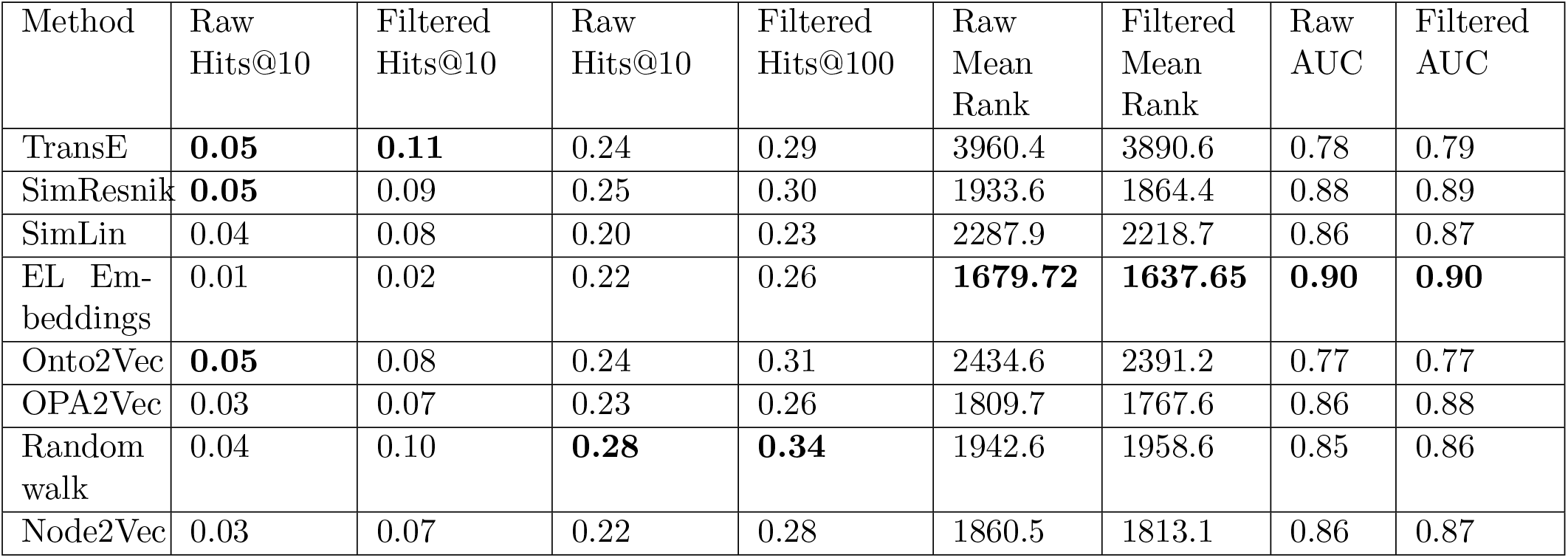
Prediction performance for human protein–protein interactions.

| Method | Raw Hits@10 | Filtered Hits@10 | Raw Hits@10 | Filtered Hits@100 | Raw Mean Rank | Filtered Mean Rank | Raw AUC | Filtered AUC |
| --- | --- | --- | --- | --- | --- | --- | --- | --- |
| TransE | **0.05** | **0.11** | 0.24 | 0.29 | 3960.4 | 3890.6 | 0.78 | 0.79 |
| SimResnik | **0.05** | 0.09 | 0.25 | 0.30 | 1933.6 | 1864.4 | 0.88 | 0.89 |
| SimLin | 0.04 | 0.08 | 0.20 | 0.23 | 2287.9 | 2218.7 | 0.86 | 0.87 |
| EL Em-beddings | 0.01 | 0.02 | 0.22 | 0.26 | **1679.72** | **1637.65** | **0.90** | **0.90** |
| Onto2Vec | **0.05** | 0.08 | 0.24 | 0.31 | 2434.6 | 2391.2 | 0.77 | 0.77 |
| OPA2Vec | 0.03 | 0.07 | 0.23 | 0.26 | 1809.7 | 1767.6 | 0.86 | 0.88 |
| Random walk | 0.04 | 0.10 | **0.28** | **0.34** | 1942.6 | 1958.6 | 0.85 | 0.86 |
| Node2Vec | 0.03 | 0.07 | 0.22 | 0.28 | 1860.5 | 1813.1 | 0.86 | 0.87 |

Overall, while our results are by no means a comprehensive evaluation and are limited to the task of predicting PPIs, we can obtain some information from our experiments. Traditional semantic similarity measures, in particular Resnik’s measure [52], performs well across many evaluations, in particular in recall at the first ranks, and often has better performance than ontology embedding methods; this property is also of importance in other applications where Resnik’s measure performs better than most other measures [30, 57]. Moreover, exploiting more of the axioms generally yields better results as can be seen when comparing EL Embeddings with most other methods. Furthermore, exploiting longer, or more indirect, relations, either through random walks or through utilizing the semantics, usually improves results over methods that are based on local properties or simply adjacency.

One experiment we did not perform here is to use the embeddings as part of a supervised machine learning model to predict the associations; such an approach has the potential to drastically improve the predictive performance results [30, 31], depending on the chosen machine learning model [23]. While we do not report on the use of embeddings for supervised learning here, we include an example of using the ontology embeddings in a Siamese neural network within the executable notebooks we provide.

## 7 Limitations and future work

Machine learning using ontologies involves a set of emerging techniques that have their roots in computer science and major applications in the life sciences where a large amount of ontologies have been developed and are applied to characterize data. Currently, several methods that allow background knowledge to be used by machine learning models are based on knowledge graphs and graph embeddings, and while these methods can be very successful, they lack the ability to utilize the model-theoretic semantics underlying ontologies. Ontologies, and representation artifacts based on similar formalisms, have the ability to represent more complex forms of knowledge, including using quantifiers, intersection, negation, and the ability to represent inconsistent knowledge. Strong negation, for example, is crucial in constraining search and cannot be substituted with the limited form of negation that is sometimes applied in knowledge graphs (i.e., the closed world assumption). However, while ontologies are able to express strong negation and other complex facts or rules, most ontology embedding methods are not yet able to adequately utilize them. Most syntactic and graph-based approaches do not interpret negation as constraints, or use any of the semantics associated with it, and can therefore not use negation to restrict search; and while model-based embeddings can utilize negation as part of the embedding they do not interact with the similarity measures or machine learning models that utilize them.

Several approaches aim to systematically integrate symbolic representations and machine learning are not yet widely applied to life science data and knowledge. Neuro-symbolic systems and neuro-symbolic integration [82, 83] provide a framework in which machine learning is integrated with symbolic representations; in the neuro-symbolic cycle, deductive inference is applied on the symbolic representations; embeddings project these representations into some space where they can be combined with data and where machine learning and optimization methods can be applied; and a knowledge extraction process maps the results back into the symbolic space. How to implement either of these projections is an active research area several of which we have reviewed here, and neuro-symbolic systems will put them together into a single framework. There is also recent interest in implementing the entire neuro-symbolic cycle, for example in vision [84]; however, with the rich set of formalized knowledge bases and the large amounts of data produced in the life sciences, we expect these systems to have major impact on how AI is applied in biology and biomedicine in the future.

Approaches to improve learning with ontologies while preserving and exploiting their semantics do not only include investigating embeddings into vector spaces (which, arguably, are mainly inspired by the needs of modern machine learning systems) but also approaches based on formal languages and logic, including Markov logic [85] and probabilistic inference [86]. Similarly, for extracting knowledge from data, new paradigms such as ‘reinforcement learning as inference’ [87] are increasingly being applied to generate explanations and representations that can be verified for consistency with background knowledge [88–90].

One main limitation of all the approaches we discussed here is their inability to consider quantitative information or data. In all cases, ontologies are used to model qualitative information and then possibly combined with other quantitative information after an embedding is generated; methods that can jointly learn on ontologies and quantitative information mapped to them include graph neural networks which will likely see increasing adoption in the coming years.

